# A region-resolved proteomic map of the human brain enabled by high-throughput proteomics

**DOI:** 10.1101/2023.06.05.543676

**Authors:** Johanna Tüshaus, Amirhossein Sakhteman, Severin Lechner, Matthew The, Eike Mucha, Christoph Krisp, Jürgen Schlegel, Claire Delbridge, Bernhard Kuster

## Abstract

Substantial efforts are underway that aim to deepen our understanding of human brain morphology, structure and function using high-resolution imaging as well has high-content molecular profiling technologies. The current work adds to these efforts by providing a comprehensive and quantitative protein expression map of 13 anatomically distinct brain regions covering more than 10,000 proteins. This was enabled by the optimization, characterization and implementation of a high-sensitivity and high-throughput micro-flow liquid chromatography timsTOF tandem mass spectrometry system (LC-MS/MS) capable of analyzing >2,000 consecutive samples prepared from formalin fixed paraffin embedded (FFPE) material. Analysis of this proteomic resource highlighted e.g. brain region-enriched protein expression patterns and functional protein classes, protein localization differences between brain regions and individual protein markers for specific brain regions. To facilitate access to and ease further mining of the data by the scientific community, all data can be explored online in a purpose-built Shiny App (https://brain-region-atlas.proteomics.ls.tum.de).

## Introduction

Around 86 Billion neurons along with a similar number of glial cells make up the most complex organ in the human body - the brain (Azevedo *et al*, 2009). The functions of the brain are highly diverse and include the control of motion, the processing of sensory information, learning and memory formation to name a few. To fulfill these complex tasks, the brain exhibits a highly-organized substructure of anatomically distinct, but well-connected brain regions. Tremendous efforts have been expended to disentangle the molecular differences of the brain regions as well as mapping their connectivity. Consortia such as the Human Protein Atlas (HPA) (Sjöstedt *et al*, 2020), the PsychENCODE project (Akbarian *et al*, 2015), the Allen brain project (Lein *et al*, 2007) and the Human brain project (HBP) (Amunts *et al*, 2019) published detailed atlases of the human brain visualizing molecular differences in a spatial dimension based on RNA-seq, MRI, in-situ hybridization and (immuno-) histochemistry data. However, a comprehensive proteomic profile of the brain regions has been lacking, arguably a substantial gap as proteins are the major functional executors of cellular processes and are the targets of almost all neurological drugs. The few proteomic studies performed so far were either focused on the mouse brain (Distler *et al*, 2020; Sharma *et al*, 2015), used pooled human tissue samples (Carlyle *et al*, 2017), were confined to one specific brain region (Guo *et al*, 2022) or suffered from limited proteome coverage (Biswas *et al*, 2021; Carlyle *et al*., 2017; Melliou *et al*, 2022). In part, this may be due to the technical hurdles involved in performing deep proteome profiling at scale. While tremendous technical improvements have been made over the years, scaling the technology to the analysis of large numbers of samples has only recently come into focus. One way of achieving greater throughput while maintaining high data quality is the use of higher chromatographic flow rates (Bian *et al*, 2021a; Bian *et al*, 2021b; Bian *et al*, 2020; Messner *et al*, 2021). We and others have demonstrated that this enables the analysis of thousands of proteomes at a moderate loss of sensitivity. To take full advantage of such improvements in peptide separation technology, mass spectrometers capable of generating data at a very rapid rate and maintaining sensitivity at the same time are required. This has, for instance, been achieved by combining trapped ion mobility and time-of-flight mass spectrometry (timsTOF), to enable the parallel accumulation and serial fragmentation (PASEF) of peptides, leading to data acquisition rates of up to 150 Hz and highly efficient utilization of the available peptide ions (Meier *et al*, 2015; Meier *et al*, 2018; Meier *et al*, 2021).

Here, we report on the coupling of micro-flow liquid chromatography (LC) to a timsTOF mass spectrometer to combine the assets of rapid and high resolution peptide chromatography with rapid and high sensitivity mass spectrometry (LC-MS/MS). We characterize the performance of this system by analyzing diverse biological samples including as human cell lines, plasma, CSF and formalin-fixed paraffin embedded (FFPE) tissue. In addition, we present an optimized end to end workflow ready for large-scale human FFPE brain proteome analysis and demonstrate its robustness by profiling 13 regions of the human brain to a depth of ∼10,000 proteins each and representing a total of >2,000 individual LC-MS/MS measurements. The resultant molecular resource, which is publically available online, constitutes the most comprehensive map of a human brain proteome to date and systematic data evaluation revealed distinct proteomic signatures of each brain region including new marker proteins.

## Results

### Optimization of a micro-flow LC timsTOF MS/MS setup

In order to establish a LC-MS/MS setup fulfilling the aforementioned requirements of sensitivity, robustness and speed, we coupled a micro-flow LC system to a Bruker timsTOF mass spectrometer equipped with a VIP-HESI ion source (Fig. 1A, B). Aiming for deep proteome coverage while restricting overall analysis time, we optimized the setup in a systematic manner including sample preparation, chromatographic, ion source and MS instrument parameters (see M&M for details).

**Figure 1.**
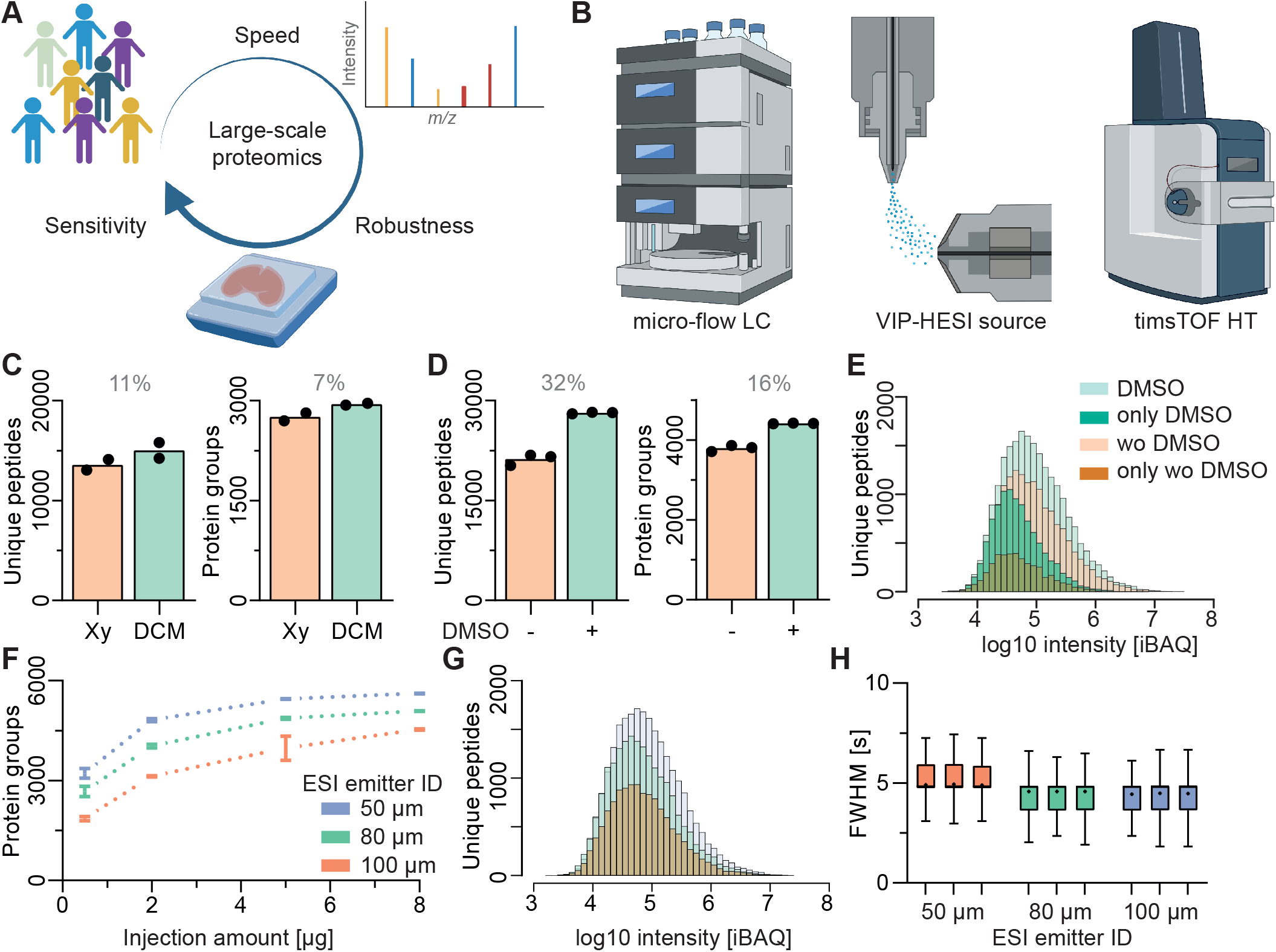
A) Schematic representation of the study aims: to establish a sensitive, robust and rapid LC-MS/MS setup able to support large-scale proteomic studies. B) Coupling a micro-flow liquid chromatography system via a VIP-HESI ion source to a timsTOF mass spectrometer. C) Bar graph indicating the number of unique peptides and protein groups identified from FFPE human brain samples using xylene (Xy) deparaffinization with or without an additional overnight delipidation step using dichloromethane (DCM) (N=2). D) Bar graph showing the number of unique peptides identified from a HeLa cell line digest in the presence or absence of 3% DMSO in LC solvents using a 30 min LC gradient and 2 µg injected digest (N=3). E) iBAQ intensity distribution of peptides identified in panel D). F) Number of protein groups identified using a 30 min LC gradient as a function of the amount of injected HeLa digest and using electrospray emitters of 50, 80 or 100 µm inner diameter (ID) (N=3). G) iBAQ intensity distribution of peptides identified in panel F) using 2 µg HeLa cell line digest loading. H) Box plots showing the chromatographic peak width distribution (full width at half maximum, FWHM) of peptides in panel G) (N=3).

First, we adapted the SP3 sample preparation approach (Müller et al, 2020) for FFPE tissue and cerebrospinal fluid (CSF) to a 96-well format using a liquid handling platform. Protein extraction from FFPE tissue was aided by sonication using the Adaptive Focused Acoustics (AFA) technology (Green *et al*, 2014; Marchione *et al*, 2020). We also added a delipidation step using dichloromethane (DCM) to the brain tissue workflow akin to methods used for tissue clearing (Molbay et al, 2021). This improved chromatographic stability and enhanced protein identification by 7% at the same time (Fig. 1C).

Second, 3% DMSO was added to LC solvents to boost electrospray ionization (ESI) efficiency (Hahne et al, 2013). This had a substantial effect on performance as it increased peptide and protein group identifications by 32% and 16% respectively using a standard 30 min LC gradient (Fig. 1D). DMSO led to an overall improvement in peptide intensity and the peptides gained in the presence of DMSO were generally of low abundance (Fig. 1E; Sup. Fig. 1A). We note that the addition of DMSO neither negatively affect the performance of the chromatographic system nor the mass spectrometer even after months of operation and thousands of sample injections. DMSO did also not change the ion mobility characteristics of analyzed peptides (Sup. Fig. 1B).

Third, the ion source parameters were optimized including the fabrication of novel ESI emitters with smaller inner diameters (ID) of 50 and 80 µm compared to the standard emitter (100 µm ID) to improve ESI efficiency. Smaller ID emitters also improve ESI stability and ion desolvation characteristics at lower liquid flow rates (Covey *et al*, 2009). As anticipated, the 50 µm emitter outperformed the wider bore emitters irrespective of the amount of material analyzed in terms of protein and peptide identifications. This was the result of a 3-fold increase in peptide intensity as well as slightly narrower chromatographic peaks widths at half maximum (FWHM) (Fig. 1E, F, H Sup. Fig. 1C, D).

Fourth, MS data acquisition parameters were optimized to take advantage of the narrow LC peak widths provided by the micro-flow LC separations. Adjustment of the threshold parameters for MS2 scheduling resulted in a gain of 3% on protein level (Sup. Fig. 1E). Reducing the time spent per MS2 scan from the standard 4.4 ms to 3.2 ms enabled an average scheduling of 18 instead of 15 precursor ions per 100 ms ramp (Sup. Fig. 1F, G). Increasing the collision energy improved protein identification by another 3% compared to standard settings and also led to a higher fraction of MS2 spectra that lead to a peptide identification (from 62 to 68%) (Sup. Fig. 1H, I). Last, we compared the timsTOF Pro2 to the timsTOF HT, the latter containing a tims-analyzer and detector designed for higher ion capacity (Sup. Fig. 1.2). At sample loadings of up to 2 µg, the timsTOF Pro2 outperformed the timsTOF HT in terms of sensitivity. However, higher sample loads resulted in a reduced number of tryptic but an increased number of semi-tryptic peptides (19%). This was likely due to overloading of the timsTOF Pro2, in turn, leading to peptide fragmentation inside the tims-analyzer. The timsTOF HT showed no such effects independent of the peptide load and outperformed the timsTOF Pro2 at loadings of more than 2 µg peptide. The following results were all obtained using the timsTOF HT.

### Evaluation of achievable proteome coverage at different levels of sample throughput

To evaluate what proteome coverage can be achieved for different levels of sample throughput, we analyzed different amounts of HeLa cell line digests using data dependent (DDA, MaxQuant) and data independent (DIA, Spectronaut 17) MS methods that would allow the analysis of between 24 and 192 samples per day (60 min respectively 7.5 min total time from injection to injection including all overhead times for sample loading, column re-equilibration etc.). For the highest throughput method, >3,100 and >5,000 proteins could be identified by using DDA or DIA, respectively and the corresponding figures for the lowest throughput method (60 min) tested were >6,600 and >8,000 proteins (Fig. 2A, Sup. Fig. 2A, B). Exemplified by data collected with the 30 min method, DDA protein identification results were almost completely contained in the DIA results (Fig. 2B). In addition, most proteins identified by DDA showed a quantitative precision of below 10%. DIA quantified a similar number of proteins with the same precision and quantified >2,500 more at still acceptable levels (up to 20% CV). (Fig. 2C, Sup. Fig. 2C).

**Figure 2.**
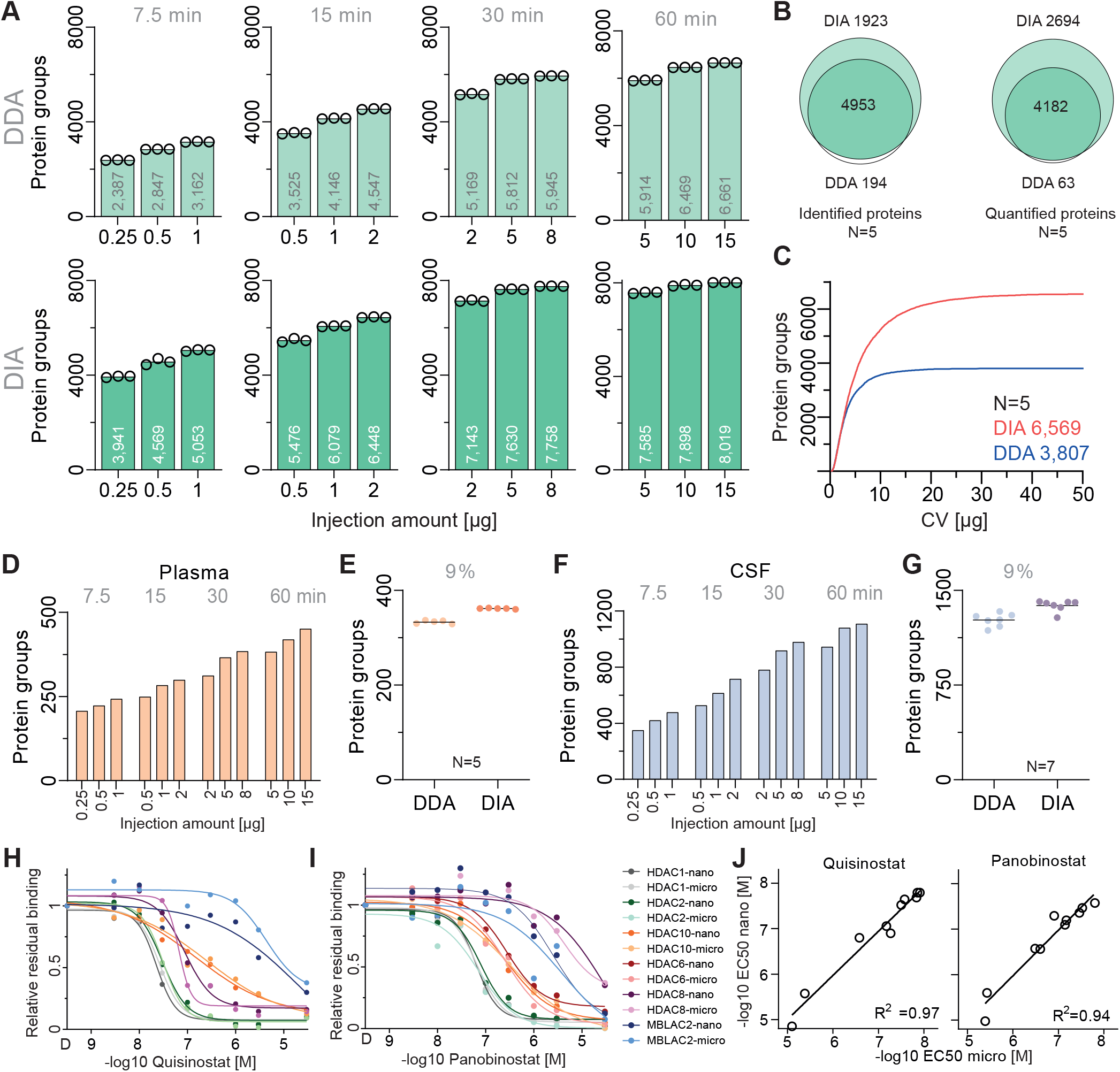
A) Number of identified proteins analysed by data dependent acquisition (DDA) (upper panels) or data independent acquisition (DIA) (lower panels) analysed by different LC gradients and as a function of the amount of HeLa digest injected (N=3). Only proteins identified by at least 2 unique peptides were considered. B) Venn diagram indicating the overlap between the number of peptides identified or quantified by DDA and DIA runs (2 µg and 30 min gradient data) (N=5). C) Cumulative density plot of the number of proteins as a function of quantitative precision (coefficient of variation (CV)) using protein LFQ intensities from data of panel B) (N=5). D) Same as panel A) but for neat digested human plasma (DDA). E) Number of proteins identified from neat human plasma digest (5 human individuals) injecting 5 µg digest and analysed using a 30 min LC gradient in DDA and DIA mode. F) Same as panel D) but for human cerebrospinal fluid (CSF). G) Same as panel E) but for human CSF. H) Dose–response binding curves of targets of the HDAC inhibitor Quisinostat obtained by competition binding assays to HDAC beads and analyzed by micro-flow timsTOF LC-MS/MS (light colors) or nano-flow Orbitrap MS/MS (dark colors). The latter data was reproduced from (Lechner *et al*., 2022). I) Same as panel H but for Panobinostat. J) Correlation analysis of -log 10 EC 50 values of dose-response curves shown in panels H) and I). R, Pearson correlation coefficient.

We next extended the evaluation to human body fluids notably plasma and CSF. In plasma, proteome coverage ranged from 243 (7.5 min) to 451 (60 min) proteins groups in single-shot DDA mode (Fig. 2D, Sup. Fig. 2.2A). Analysis of five individual plasma samples (5 ug each; 30 min method) resulted, on average, in 333 and 362 proteins using DDA and DIA, respectively (Fig. 2E, Sup. Fig. 2.2B). Notably, 45 (DDA) and 47 (DIA) of the 49 FDA-approved biomarkers previously identified by (Geyer *et al*, 2016) were covered in the dataset with more than 2 unique peptides. At the same level of sample throughput, 1,283 and 1,375 proteins were, on average, identified in CSF samples in DDA and DIA mode, respectively (Fig. 2F, G, Sup. Fig. 2.2B). Comparable CSF proteome coverage was recently reported using nano-flow MS/MS and LC gradients of 45 - 80 min (Bader *et al*, 2020; Karayel *et al*, 2022).

In a third application, we used the 30 min DDA method to profile the targets of HDAC-inhibitors using a competition binding assay previously established by the authors requiring 110 min of total nano-flow LC-MS/MS time (Lechner *et al*, 2022). Exemplified by the three HDAC inhibitors Quisinostat, TSA and Panobinostat, the micro-flow timsTOF setup provided very similar data quality in terms of detection of target proteins as well as drug:target interaction strength but in a fraction of total analysis time (Fig 2H, I, J, Sup 2.2C, D).

### Deep proteomic profiling of 13 formalin-preserved human brain regions

With an optimized, high-throughput-capable micro-flow LC timsTOF HT system at hand, we set out to establish a region-resolved proteomic atlas of the human brain. Thirteen brain regions were collected from one formalin-preserved postmortem human brain (Nucleus accumbens, Substantia nigra, Olives, Thalamus, Red nucleus, Hippocampus, Putamen, Claustrum, Caudate nucleus, Cerebellum and Cerebral cortex (grey & white matter of the letter two)). From each brain region, three tissue cubes (∼5x5x5 mm) were collected and independently processed (two in case of olives). Following protein extraction and digestion, each of the 38 resulting peptide samples was separated into 48 fractions using high-pH reversed phase chromatography and analyzed by micro-flow LC-MS/MS using the 30 min method described above (Fig 3A). Including control samples (see below), this led to collecting data from 2,271 LC-MS/MS runs, all using the same online C18 column and the same 50 µm ESI emitter. To control for stable technical performance of the micro-flow LC-MS/MS setup, full proteome HeLa digests were analyzed between tissue samples (N=21). Almost all proteins (98% of 2,845) and 76% of all peptides (10,488) showed CV values of below 20% (Fig 3B). The median CV was 7% on protein and 12% on peptide level. For the brain regions the respective median protein CVs were somewhat higher (10-20%). This is due to the detection of a far greater number of proteins in each sample (>9,000 proteins), the additional step of peptide fractionation, and the biological diversity within a brain region (Fig. 3C). To measure LC performance stability, synthetic (PROCAL) peptides, designed to span the entire peptide elution spectrum across an LC gradient (Zolg *et al*, 2017), were run either alone or spiked into each tissue sample (N=1,968). The CVs of retention times were very small (1%) and nearly identical when run alone or as spike-in (Fig. 3D, Sup. Fig. 3A).

**Figure 3.**
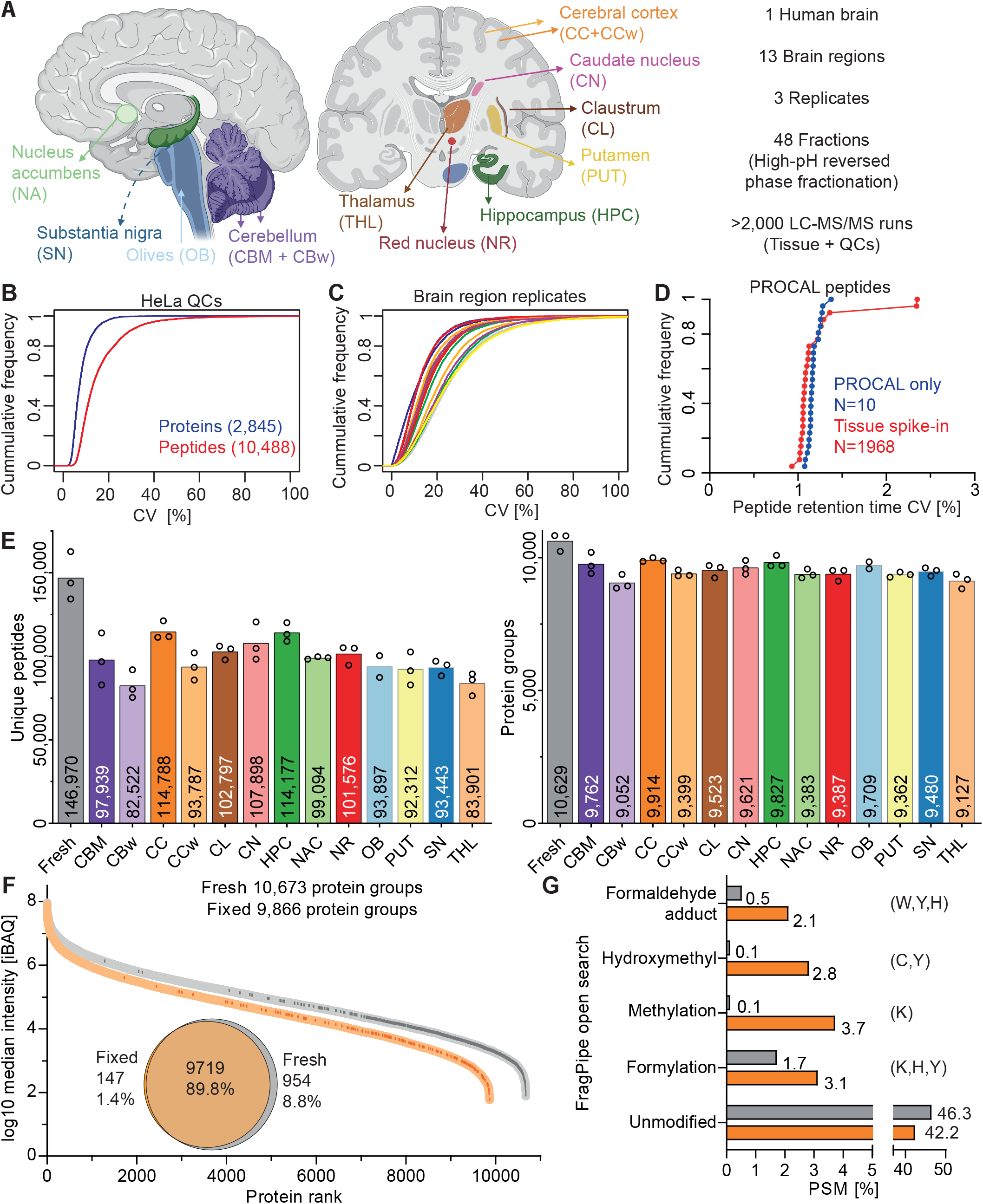
A) Illustration of the brain regions included in the proteomic atlas and numerical summary of the scope of the study. B) Cumulative density plot of the number of identified proteins and peptides as a function of quantitative precision (coefficient of variation (CV)) using protein LFQ intensities (2 µg HeLa peptides, 30 min LC gradients, N=21). C) Same as panel B) but for brain proteins from the different regions analysed in this study. D) Same as panel B but for chromatographic retention time precision of synthetic peptides (PROCAL) run as LC quality controls throughout the project either alone (blue, N=10) or spiked into fractionated brain region digests (red, N=1968). E) Bar graph showing the number of unique peptides and protein groups identified from each brain brain region (Cerebellum (CBM), white matter cerebellum (CBw), cerebral cortex (CC), white matter cerebral cortex (CCw), claustrum (CL), caudate nucleus (CN), hippocampus (HPC), nucleus accumbens (NAC), red nucleus (NR), olives (OB), putamen (PUT) substantia nigra (SN), thalamus (THL)). F) Rank plot of proteins from either fresh-frozen (grey) or formalin-preserved (orange) cortex sorted by iBAQ rank. Dark marks within the rank line indicate the abundance of proteins only found in one of the two samples. The Venn diagram shows the overlap at protein group level. G) Results of an open modification search using FragPipe comparing the most common chemical modifications attributed to formaldehyde fixation (PSM, peptide spectrum match).

Deep proteome coverage was achieved of all brain regions with, on average, 9,498 proteins and 98,434 peptides identified in each tissue (Fig. 3E) and a total of 11,325 proteins across all regions. As a further indicator of data quality and achievable proteome coverage, three replicates of fresh frozen tissue of the cerebral cortex of a different postmortem human brain was analyzed in the same manner to estimate the losses caused by the fixation process. Proteome coverage of the fresh frozen CC material (average of 10,629 protein groups and 146,970 peptides per replicate) was ∼7% deeper than that of FFPE CC material and the difference was much more pronounced at peptide level (28%). Comparing the intensities of the two samples showed that both proteomes span about six orders of magnitude in protein expression (Fig 3F). Almost all proteins (98.6%) detected in the fixed tissue were contained in the data of the fresh frozen sample but had systematically lower intensities. Most proteins exclusively detected in the fresh frozen tissue populate the lower abundance range (87% of quartile 4) which was more pronounced than for proteins exclusively detected in the fixed material (Fig 3F, Sup 3B). iBAQ ratios of proteins identified in common between fresh and fixed tissue revealed an abundance-dependent pattern indicating some low abundance proteins in fresh tissue to be overrepresented in fixed tissue (Sup. Fig. 3C). According to functional clustering analysis, nuclear proteins related to transcription were overrepresented in the top 10% of proteins high in the fresh relative to fixed tissue. In contrast, the bottom 10 % of proteins of low abundance in fresh relative to fixed tissue were associated with secretion and the extracellular matrix. An open modification search using FragPipe (Kong *et al*, 2017; Yu *et al*, 2020a; Yu *et al*, 2020b) returned 46% of all peptide spectrum matches (PSMs) in fresh tissue as unmodified, compared to 42% in the fixed tissue. In contrast, many more modifications that can be rationalized by the use of formaldehyde in the fixation process (Metz *et al*, 2004) were indeed found in the fixed tissue (Fig. 4D). The above data generally indicates that formaldehyde crosslinking cannot be fully reversed leading to substantial loss of peptides that can be recovered or identified from FFPE material. At the same time, the data shows that certain proteins can be more efficiently extracted from FFPE than fresh tissue.

**Figure 4.**
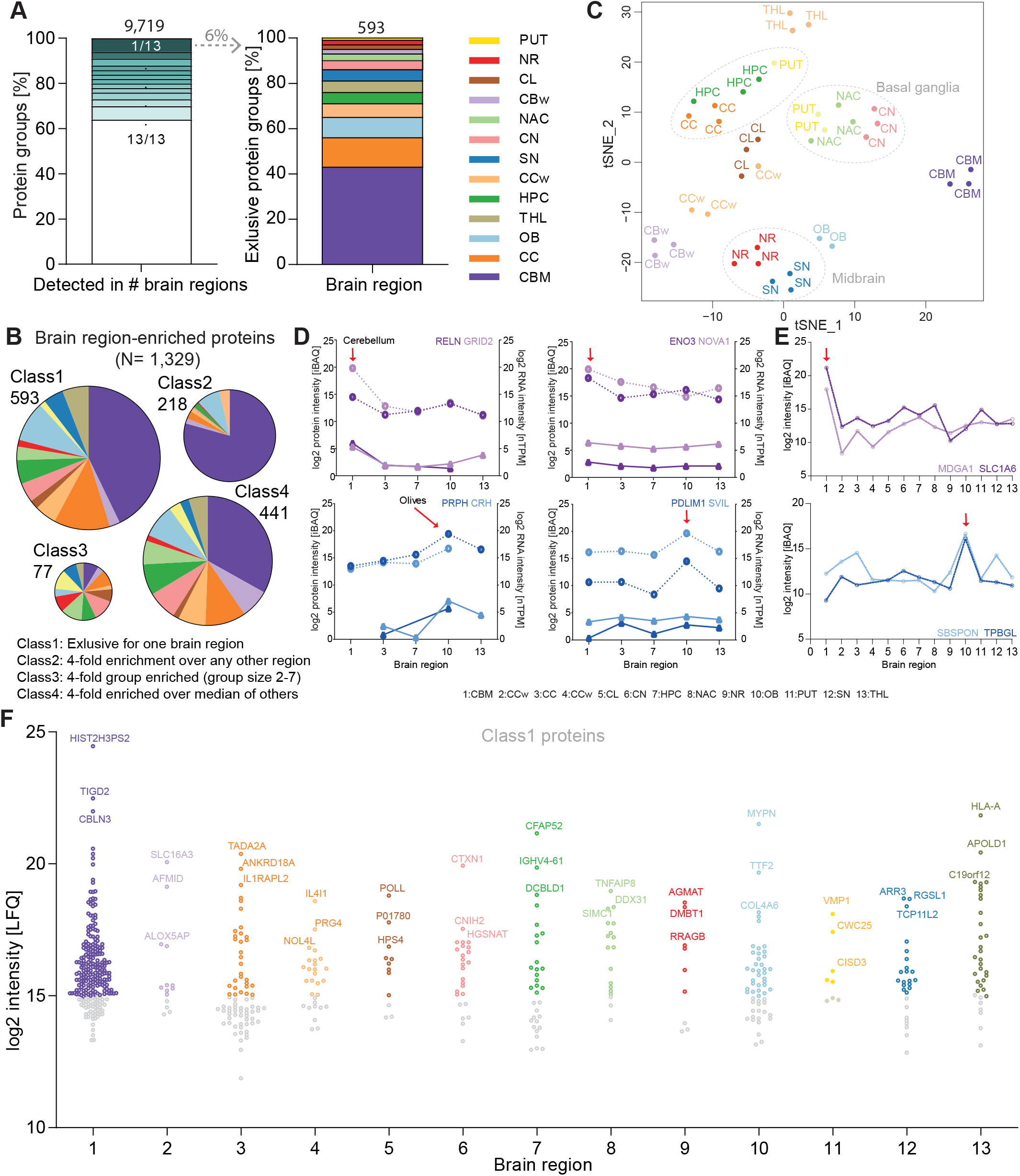
A) Percentage of proteins detected in at least two biological replicates and their distribution over 13 brain regions (left panel). Right panel: distribution of the 593 proteins exclusively identified in one brain region. B) Pie charts showing the distribution of proteins in four enrichment classes across the brain regions. C) tSNE plot of all replicate proteomes from all brain regions. Illustrating their proximity by considering all proteins. Proximity of basal ganglia (NAC, CN, PUT) and midbrain (NR, SN) samples are highlighted by dotted lines. D) Comparison of protein and mRNA levels of example proteins in five brain regions. The mRNA data was taken from the human protein atlas (HPA) (Sjöstedt *et al*., 2020). Red arrows point to brain regions in which the protein is statistically significantly higher expressed as in other brain regions. E) Protein expression profiles of four example proteins across all 13 brain regions. Red arrows as in panel D). F) Swarm plot showing Class1 proteins (i.e. proteins exclusively identified in on region) sorted by LFQ intensity. Low abundant proteins (log2 LFQ < 15) are shown in grey.

### Analysis of the brain region-resolved human proteome atlas

The proteome atlas generated in this study offers a multitude of analyses only a few of which can be highlighted in the following. According to our atlas, 64% of all proteins were detected in all 13 brain regions and 6% were exclusive to a single one. Most of the exclusive proteins were found in the cerebellum (CBM), followed by cerebral cortex (CC), olives (OB) and hippocampus (HPC) (Fig. 4A, Sup. Fig. 4.1A, B). Interestingly, as many as 5,750 of the 9,719 proteins in the atlas were significantly differentially expressed between brain regions (ANOVA, followed by pairwise T-tests for multiple comparisons of independent brain regions) (Sup. Table 1). Following a concept adopted from the Human Protein Atlas (HPA) (Sjöstedt *et al*., 2020), we defined four regional protein enrichment classifications. Class 1 proteins (593) were only detected in a single brain region, Class 2 (218) includes proteins that are at least 4-fold enriched in one brain region compared to all others, Class 3 (77) are so-called group-enriched proteins (i.e. at least 4-fold enriched in 2-7 brain regions) and Class 4 (441) comprises regionally enhanced proteins (at least 4-fold enriched in one region over the average of all other regions) (Fig. 4B, Sup. Fig. 4.1C). The rational for the 4-fold enrichment was derived from modeling the quantitative variance from the three replicates of PUT (the region with the highest variance in the dataset). Using this criterion, protein expression differences between brain regions can be detected with 99.998% confidence (Fig. 3C, Sup. Fig. 4D). By applying this extremely stringent cut-off, we aimed to focus attention on likely biologically meaningful protein expression differences between regions. In total, 1,329 proteins of 9,719 (14%) fulfilled this criterion in any of the above four classes and are, from here on, referred to as brain region-enriched proteins (Sup. Table 1).

A closer look at the brain region distribution within the different enrichment classes, highlighted the predominance of CBM within class 1, 2 and 4. In contrast, class 3, which includes the group-enriched proteins, revealed a more even distribution between the brain regions. These include proteins such as BCL11B, CHAT and SLC10A4, enriched in putamen (PUT), caudate nucleus (CN) as well as nucleus accumbens (NAC). These three regions are all part of the basal ganglia. The two brain regions of the midbrain, substantia nigra (SN) and red nucleus (RN), also share group-enriched proteins such as RTL1 and IL17RA. tSNE analysis as well as hierarchical clustering of all the data confirmed that anatomically close brain regions often share protein expression patterns (Fig. 4C, Sup. Fig. 4.3A, 4.4).

Next, we compared the proteomic and transcriptomic profiles of five brain regions for which mRNA data was available from the HPA project (Sjöstedt *et al*., 2020). As observed many times before, the overall correlation between mRNA and protein levels was low (Sup. Fig. 4.2) (Carlyle *et al*., 2017; Wang *et al*, 2019). Similarly, while there are many cases where the trends in protein and mRNA levels were similar between brain regions (Fig. 4D left panels), there are also many cases for which mRNA levels were more stable than protein levels (Fig. 4D right panels). These observations yet again underscore the importance of measuring protein expression directly rather than relying on mRNA levels as a proxy, particularly when it comes to the identification of markers for certain brain regions. Class 1 proteins are the most likely source for such markers and many well-known cases were found within this class including MDGA1 and SLC1A6 in cerebellum as well as SBSPON and TPBGL in the medulla oblongata, the larger sub-region of the olives (Fig 4E). Dozens of further candidates were identified for most brain regions and many of these are of high abundance ruling out the possibility that these are technical artefacts (Fig. 4F, Sup. Fig. 4.3 A, B, C). Examples include the GPCR-associated signaling protein ARR3 in substantia nigra or the small (9 kDa) but poorly characterized protein CTXN1 in caudate nucleus.

A previously published transcriptome analyzes of the major brain cell types (neurons, microglia, astrocytes, oligodendrocytes) by the HPA has defined cell type-specific signatures based on mRNA expression and many of these have also been detected in the current study (Fig 5A). Of the 1,329 brain region-enriched proteins classified in the current work, 824 (62%) also fall into the cell type-specific mRNA HPA classification. These 824 regionally-enriched and supposedly cell type-specific proteins split nearly evenly between neurons (46%) and glial cells (54% encompassing microglia, astrocytes, and oligodendrocytes (Sup. Table 1). About 85% of the glial mRNA-based signature proteins were detected in all brain regions and in similar quantities, exemplified by the oligodendrocyte marker MBP, the astrocyte marker GFAP and the microglia marker RGS10. The remaining, 15% showed enrichment in at least one brain region including the oligodendrocyte protein MTUS1 in OB as well as the astrocytic proteins, NFIA and NFIB, in CBM (Fig. 5B). Similarly, the majority of the neuronal mRNA-based signature proteins (83%) were not regionally enriched including the neuronal marker proteins CD200, SYNJ2BP and SPTBN4 (Fig. 5C) but about 17% were, including MLIP in CC and TPBGL in OB. A closer look at the brain region distribution of the 824 region-enriched mRNA signature proteins, showed that e.g. oligodendrocyte signature proteins are particularly prevalent in cortical white matter (CCw) (Fig. 5D) with a 1.3-fold mean abundance ratio over the average of all other brain regions (Fig. 5E). This may be rationalized by the characteristic architecture of the white matter which mainly consists of neuronal axons that are encapsulated by oligodendrocytes. In contrast, neuronal signature proteins were most prevalent in cortex (1.4-fold) (Sup. Fig. 5.1B) while the majority of synaptic signature proteins (as classified by UniProt) were detected in CC followed by HPC and CN (1.7, 1.6, 1.5-fold respectively) (Fig. 5D, F). Among the top 10 most enriched synaptic proteins in the cerebral cortex are SHISA6 and SHISA7 which control synaptic transmission in excitatory neurons as well as SYNPO and LRRC7 involved in spine architecture (Fig. 5G).

**Figure 5.**
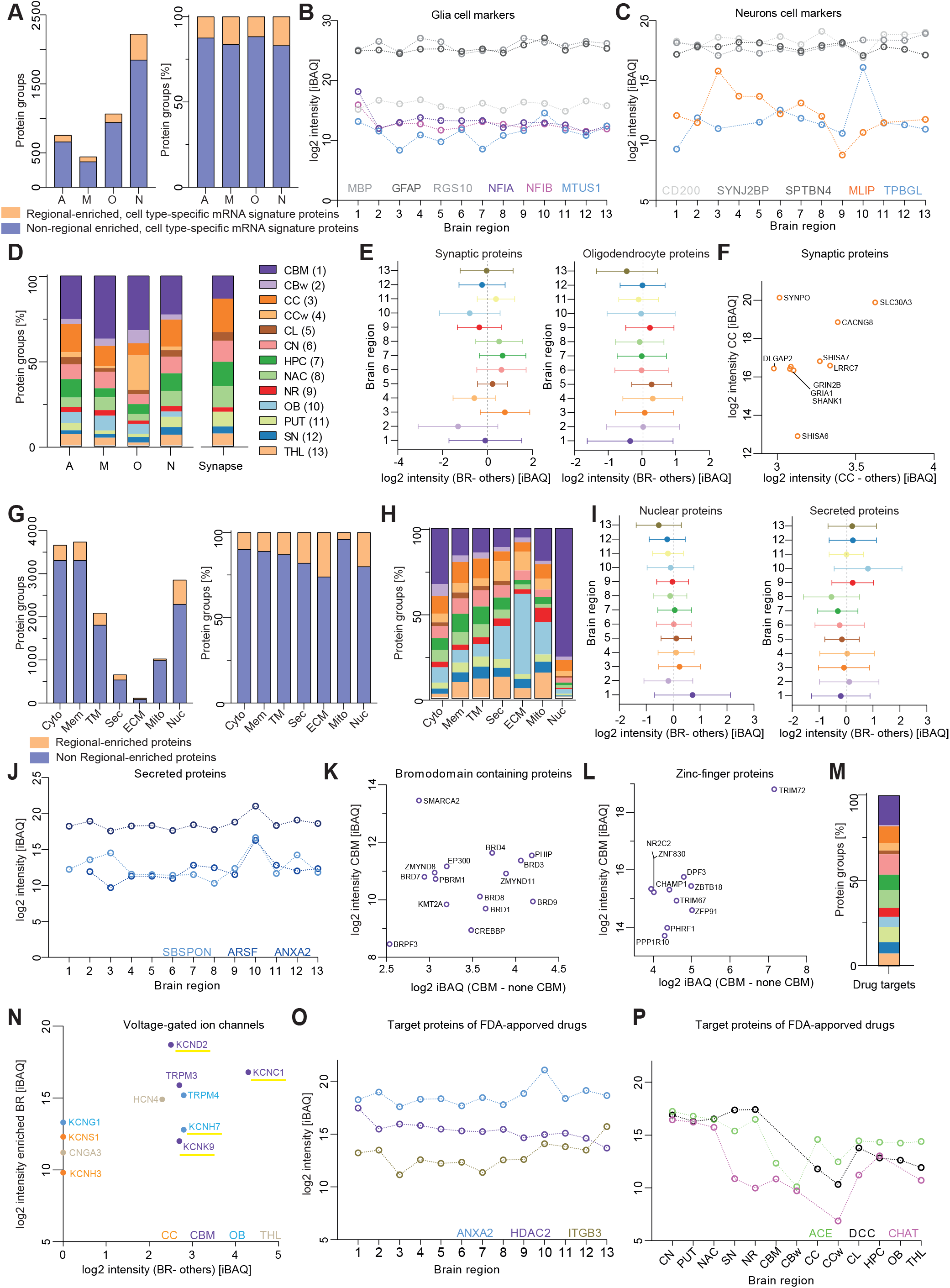
A) Bar graph showing the absolute and relative number of cell type-enhanced protein markers identified previously by the HPA project based on mRNA data (Sjöstedt *et al*., 2020) which either exhibit a regional enrichment in our proteome atlas (yellow) or not (blue). The four major brain cell types neurons (N) and the glial cell types astrocytes (A), microglia (M) and oligodendrocytes (O) are included in the analysis. B) Protein abundance profiles of cell type-enriched proteins of glial cells across the 13 brain regions. C) Same as panel B) but for neuronal cells. D) Contribution of the different brain regions to the regionally-enriched, brain cell type-enhanced proteins (yellow proteins in panel A). The regional distribution of synaptic proteins according to UniProt is shown in addition. E) Protein abundance ratios of all synaptic proteins (according to UniProt) and oligodendrocyte enhanced proteins (according to HPA) in each specific brain region over the average of all other brain regions (dotted line). F) Mean (N=3) log2 iBAQ intensities of the top 10 most abundant synaptic proteins in cerebral cortex (CC) plotted over the difference to the mean of all other brain regions. G) Absolute and relative number of proteins sorted by cellular compartment (according to UniProt) of regional enriched proteins (yellow) and non-regionally enriched proteins (blue) in the brain proteome atlas (cytoplasm (Cyto), membrane (Mem), transmembrane (TM), secreted (Sec), extracellular matrix (ECM), mitochondria (Mito), nuclear (Nuc)). H) Relative contribution of the 13 brain regions to the regionally-enriched proteins (yellow in panel G). I) Protein abundance ratios of all nuclear or secreted proteins (according to UniProt) in each specific brain region over the average of all other brain regions (dotted line). J) Examples of protein abundance profile of secreted proteins enriched in olives (OB). K) Mean (N=3) log2 iBAQ intensities of the top 15 most abundant bromodomain-containing proteins in cerebellum (CBM) plotted over the difference to the mean of all other brain regions. L) Same as panel K) but for zinc finger-containing proteins. M) Proportion of the different brain regions to the regionally-enriched proteins which are classified as drug targets of FDA-approved drugs (according to drugbank, https://go.drugbank.com/). N) Abundance of voltage-gated ion channels enriched in specific brain regions. The colour code follows the brain regions and the yellow underlining indicates proteins which are targets of FDA-approved drugs. O) Protein abundance profiles of brain region-enriched proteins that are targets of FDA-approved drugs. P) Further examples akin to panel O).

We next asked if brain region-enriched proteins exhibit differences in cellular localization (as classified by UniProt) compared to all proteins detected in this study. While the majority of all detected proteins were localized to the cytoplasm or membrane-associated, most of the brain region-enriched proteins were annotated as nuclear, followed by membrane-associated, cytoplasmic, transmembrane and secreted (Fig. 5G). Interestingly, secreted and ECM-associated proteins were highly overrepresented in OB (log2 of 1.8-fold) (Fig. 5H, I). These include the signaling cue proteins SBSPON, ARSF and ANXA2 (Fig. 5J). This observation is also reflected in the GO-term analysis of OB in which signal transmission and extracellular matrix are major functional annotations (Sup. Fig. 6). Brain region-enriched proteins of nuclear localization were predominantly over-represented in cerebellum (log2 of 1.8-fold) (Fig. 5I, J) and included 15 members of the bromodomain and 163 members of the zinc-finger families of proteins (Top 10 enriched shown in Fig. 5K, L). Again, this observation is backed by GO-term and KEGG analysis of CBM which highlights DNA- and RNA-binding, transcription and the spliceosome as major functional annotations (Sup. Fig. 6). This relative overrepresentation of nuclear proteins within CBM is likely owing to the high density of neurons with enlarged nuclei within the grey matter of the cerebellum.

Perhaps anecdotally, we detected 12 neuropeptides of which three showed a regional enrichment. The endocrine hormones GAL and PNOC belong to class 1 enriched in OB and CC, respectively. In addition, PENK, a neuropeptide involved in pain perception (an opioid mimic) had high levels in midbrain (SN, NR) and basal ganglia (CN, PUT, NAC) (Sup. Fig. 5.1C). PENK was previously shown to be involved in glutamate release within the striatum (CN, PUT).

We then matched a list of proteins targeted by FDA approved drugs (https://go.drugbank.com/) to the proteomic data and identified 470. Of these, 68 showed a brain region-specific enrichment covering all 13 brain regions (Fig. 5M). Among the 68 were 11 voltage gated ion channels (Fig. 5N), and ITGB3, ANXA2 and HDAC2 (Fig. 5O). The latter is noteworthy because only HDAC2 showed a regional enrichment in cerebellum while all other detected HDAC family members were found with similar levels in all brain regions. Similarly, 8 members of the annexin family were detected of which only ANXA2 and ANXA3 showed regional enrichment in CCw. In addition, several group enriched (Class 3) proteins classified as drug targets were over-represented in basal ganglia (ACE, DCC, CHAT) and midbrain (ACE, DDC) (Fig. 5P).

The above examples served to exemplify the types of questions one might pose to the data. In order to make the atlas easily accessible to the scientific community, we have created a web-based shiny app that enables visualizing the expression of any protein across any brain region (https://brain-region-atlas.proteomics.ls.tum.de/main_brainshinyapp/) and also offers tabular download of selected data and graphics.

## Discussion

Tremendous efforts have been expended to map regional diversity in the brain mostly by using imaging and transcriptomic approaches that all attempt to help understand the highly organized substructures and diverse functions of the human brain. The current work makes a number of valuable contributions in this context. First, we developed a scalable, yet sensitive LC-MS/MS approach that paves the way for larger-scale analysis, e.g. by eventually analysing all anatomically or functionally distinct regions of the brain or comparing brain structures of many individuals in terms of pathophysiology. Using this setup, the proteomic depth can be tuned seamlessly from 5,000 to 10,000 proteins by allocating either a few minutes or a few hours of time and requiring only single digit microgram quantities of total protein. Compared to the state of the art in micro-LC-MS/MS on Orbitrap instruments (Bian *et al*., 2020), the new setup achieves very similar proteome coverage in 75% less time and using 90% less sample. An important advantage over traditional nanoLC-MS/MS is the noteworthy robustness of the setup as the entire brain region atlas project with >2,000 consecutive samples was developed using the same LC column and ESI emitter. Second, we developed an efficient protocol for protein extraction from formalin-preserved brain material that enabled the collection of quantitative expression information for ∼10,000 proteins from each brain region. This goes far beyond what was previously reported for large-scale FFPE material studies where proteome depth was limited to ∼5,000 proteins (Bhatia *et al*, 2022; Coscia *et al*, 2020; Eckert *et al*, 2021). The fact that the data on FFPE material is nearly as deep as that for fresh-frozen holds enormous potential for the analysis of the millions of archived tissue specimen in human biobanks. Third, analysis of the 13 brain regions uncovered unique proteomic fingerprints including well-established regional marker proteins but also many new candidates. We found numerous cases where mRNA and protein profiles substantially deviated from each other underscoring the importance of measuring protein levels directly. There is also strong evidence for differences in cellular localization of proteins between brain regions as well as the expression of annotated drug targets. Forth, the web-based Shiny App along with the deposited MS raw files, protein identification and quantification data will serve the neuroscience community as a valuable data mining tool for many lines of investigations not covered in the current manuscript.

Despite the advances outlined above, many challenges remain. For instance, while we deconvoluted several substructures of e.g. the basal ganglia (caudate nucleus, putamen, nucleus accumbens) and the midbrain (substantia nigra, red nucleus), many more exist but which were not covered here. Similarly, the atlas does not cover all brain cell types, so the contribution of rare cell types such as pericytes are likely overlooked. In addition, there is no component of spatial organization yet, for instance regarding differences in protein expression within a brain region or between the hemispheres. And, last, the atlas currently constitutes a static picture of a single human brain, so it does not shed any light on differences between individuals, dynamic changes during e. g. development or the onset and progression of disease. Still, the proteomic technology presented in this study has the potential to address most of these points because its throughput and robustness scales to the analysis of >15,000 FFPE samples per year that may e.g. be deployed to mapping the brain or its regions at higher special resolution or focused parts of the brain across many biological or pathological conditions.

## Materials and Methods

### Brain tissue

The formalin fixed brain of a 56-year old Caucasian male was dissected coronally into 5 mm thick slices. After no specific diagnostic observations were made upon macroscopic inspection and the diagnostic report was completed by a neuropathologist, 5x5x5 mm cubes of both hemispheres of the following 12 brain regions were collected: Nucleus accumbens, Substantia nigra, Thalamus, Red nucleus, Hippocampus, Putamen, Claustrum, Caudate nucleus, Cerebellum and Cerebral cortex (grey & white matter of the latter two). Additionally, two cubes of the olives were collected and stored in formalin until further processing.

### Sample preparation

HeLa cell lysate and plasma samples were diluted in 8 M urea buffer (80 mM Tris-HCl, pH 7.6) based on a protocol by (Bian *et al*., 2020). A Bradford assay was used for protein concentration estimation. Proteins were denatured with 10 mM DTT for 60 min at 37°C while shaking at 700 rpm. Next, 55 mM 2- chloracetamid (CCA) was added at 37°C while shaking for 60 min at 700 rpm. Five volumes of 50 mM Tris pH 8 was added and proteins were digested over night with trypsin (1:50) (Roche) at 37°C while shaking at 800 rpm. Formic acid was added to quench the digest (1% final concentration) and the peptides were at 5000xg for 15 min. Acidified peptides were desalted using Sep-Pak columns (HeLa) according to the user manual or using stage tips (plasma) (Rappsilber *et al*, 2007).

100 µL CSF per individual was digested using the SP3 approach (Hughes *et al*, 2019). In brief, CSF was denatured with 15 mM DTT in 50 mM ammonium bicarbonate (ABC) for 30 min at 45°C while shaking at 1200 rpm, alkylated with 50 mM CAA in 50 mM ABC while incubating for 30 min at room temperature in the dark. Next, 15 mM DTT in 50 mM ABC was added. SP3 beads were washed with water and 5 µL beads (1:1) were used for protein binding in the presence of 70% ethanol. After washing with 80% ethanol, proteins were digested with trypsin (1:16) overnight at 37°C while shaking at 1200 rpm. Peptides were desalted with stage tips (Rappsilber *et al*., 2007), dried in a Speed Vac and peptide concentrations were estimated using a Nanodrop.

Brain tissue cubes were incubated overnight in xylene followed by dichloromethane (DCM), washed three times with 100% ethanol, 96% ethanol, 70% ethanol and water. Next, the tissue was ground using a tissue lyser (Qiagen, 3 min at 300 s^-1^) in 200 µL 500 mM Tris pH 9 by adding a metal ball (diameter 5 mm). Tissue lysis and decrosslinking was performed according to (Eckert *et al*., 2021) using 4% SDS, 10 mM DTT in 500 mM Tris pH 9. The samples were incubated for 90 min at 95°C while shaking at 1200 rpm. In between, after 45 min of decrosslinking, samples were transferred to a 96 AFA-TUBE TPX Plate (520291) (100 µL per well) and sonicated for 5 min using the Covaris FPPE protocol using the R230 Focused Ultrasonication Device from Covaris (peak power: 350, duty factor: 25, cycles per burst 200, average power: 87.5). After an additional 45 min of crosslinking, protein concentration was measured using the PierceTM 660 nm protein assay kit according to the manufacture manual (with 0,25g α- Cyclodextrin per 5 mL 660 nm assay solution). The pH of the lysate was adjusted to pH 7 using 8% formic acid. Next, 3x 200 µg protein lysate (pH 7) of each sample was digested according to the SP3 protocol on a Bravo handling platform (Bian *et al*., 2020; Müller *et al*, 2020). In brief, 20 µL Sera Mag A–B bead mix (1:1) was used per 200 µg total protein lysate sample and proteins were bound at a final concentration of 70% ethanol. Beads were washed with 80% ethanol and acetonitrile (ACN). Proteins were denatured with 10 mM DTT (45min, 37C) followed by alkylation with 50 mM CAA in 40 mM Tris pH 7.6, CaCl2 2mM (30min, 25C). Trypsin digestion was performed overnight (1:50). On the next day peptides were collected, beads were washed with 2% formic acid and desalted with Sep-Pak columns according to the user manual.

Off-Line High-pH Reversed-Phase Fractionation was performed using a Dionex Ultra 3000 HPLC system and a Waters XBridge BEH130 C18 column (3.5 µl 2.21 x 250 mm) based on (Bekker-Jensen *et al*, 2017). 200 µg desalted peptides were reconstituted in 25 mM ABC pH 8 and separated with a linear gradient from 5% Buffer B to 40% Buffer B within 48 min at a flow rate of 200 µl/min (Buffer A: H_2_O MS grade, Buffer B: ACN, Buffer C: 25 mM ABC pH 8 constant at 10%). Every 30 s a fraction was collected, ending up in 96 fractions which were pooled to 48 (combining fraction 1 and 49 etc.). Fractionated peptides were frozen, dried with a SpeedVac and reconstituted with 0.1% FA plus 100 fmol PROCAL peptides. 50% of each fraction (2 µg) were injected for LC-MS/MS analysis.

### Mass spectrometry

All proteomic data were acquired with a micro-flow LC coupled via a VIP HESI source to a timsTOF mass spectrometer. Liquid chromatography was performed on a Dionex UltiMate 3000 System including a WPS-3000TPL autosampler allowing direct sample pickup together with a NCS-3500RS Nano ProFlow equipped with a micro-flow selector within the flowmeter and standard nano pumps or a Vanquish Neo UHPLC-System in mirco-flow mode (Thermo Scientific). All connections were closed via NanoViper capillaries with 50 µm ID (Thermo Scientific). A 20 µL sample loop was used. Peptides were separate on a Pepmap C18 column (1 mm ID, 15 cm lengths, 2 µm particle size) at a flow rate of 50 µL/min. Binary gradients (listed below) of buffer A and B were run (A: 0.1% FA in H_2_O, B: 80% ACN 0.1%FA) including 3% DMSO, if not stated otherwise. The column oven temperature was set to 60°C. Sample loading, column washing, and equilibration was performed at maximum speed.

**Table 1:**
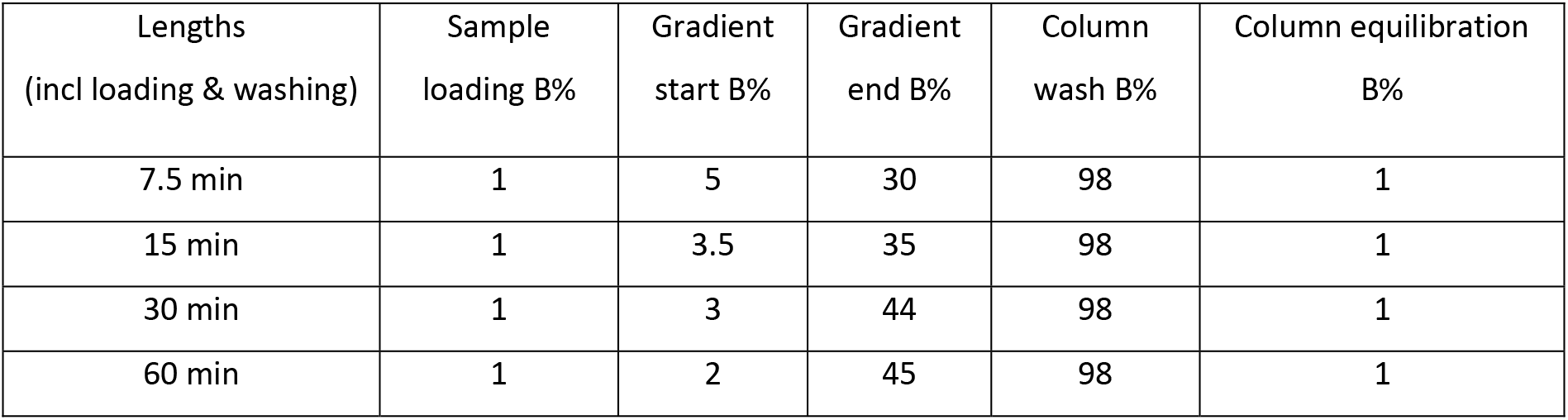
LC gradients

| Lengths<br>(incl loading & washing) | Sample<br>loading B% | Gradient<br>start B% | Gradient<br>end B% | Column<br>wash B% | Column equilibration<br>B% |
| --- | --- | --- | --- | --- | --- |
| 7.5 min | 1 | 5 | 30 | 98 | 1 |
| 15 min | 1 | 3.5 | 35 | 98 | 1 |
| 30 min | 1 | 3 | 44 | 98 | 1 |
| 60 min | 1 | 2 | 45 | 98 | 1 |

The source parameters were optimized in regard of signal stability as well as signal intensity and were kept constant over all measurements. Different prototypes of emitters (IDs: 50, 80, 100 µm) and union connections between emitter and NanoViper capillary connected to the column were tested.

**Table 2:**
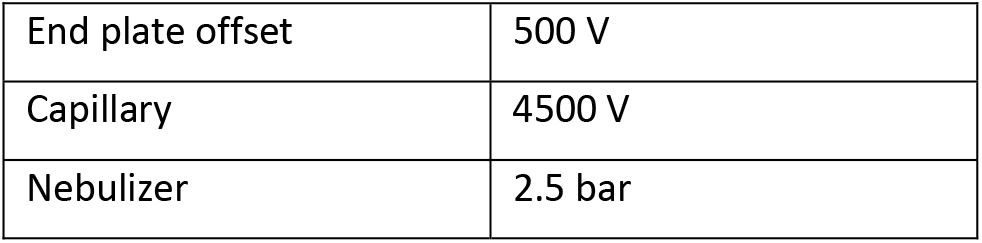
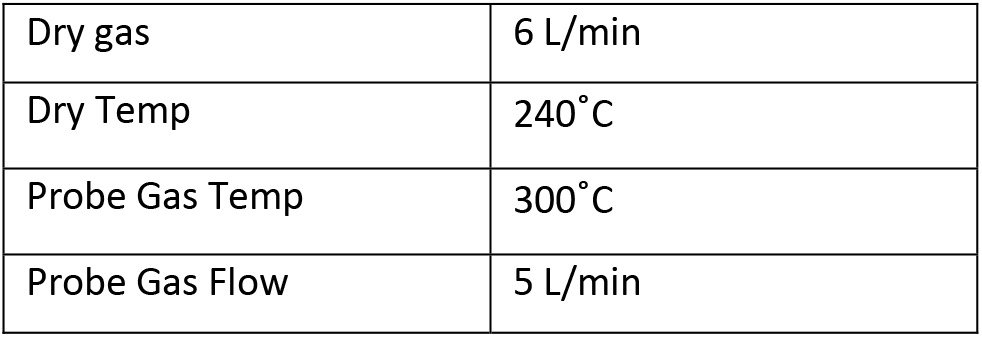
VIP HESI source settings

A timsTOF HT system was used for the majority of the measurements. Only the measurements depicted in Supplementary data 1.2 were acquired on a timsTOF Pro2 system. DDA runs were acquired in PASEF mode in a mobility range of 0.85 and 1.3 with a ramp time of 100 ms with a duty cycle of 100%. The number of ramps varied according to the gradient lengths to control for sufficient data points for quantification (8 p per peak) between 4 and 10. Collision energy settings were optimized to 59 eV at 1/K0 of 0.6 Vs/cm² to 29 at 1/K0 of 0.6 Vs/cm², target intensity to 12,000 and intensity thresholds 1,600. MS2 scheduling was fastened from standard quadrupole switching time of 1.6 ms to 1.2 ms and MS2 acquisition time from 2.75 ms to 2 ms. DIA runs were acquired in PASEF mode in a mobility range of 0.64 to 1.45 with a ramp time of 100 ms with a duty cycle of 100%. Advanced collision energy settings were enabled. The DIA window scheme followed a 3x 8 pattern of 25 m/z widths covering the whole mobility range as following:

**Table 3.** DIA window scheme

| #MS Type | Cycle Id | Start IM [1/K0] | End IM [1/K0] | Start Mass [m/z] | End Mass [m/z] |
| --- | --- | --- | --- | --- | --- |
| MS1 | 0 | - | - | - | - |
| PASEF | 1 | 1.01 | 1.37 | 800 | 825 |
| PASEF | 1 | 0.83 | 1.01 | 600 | 625 |
| PASEF | 1 | 0.64 | 0.83 | 400 | 425 |
| PASEF | 2 | 1.04 | 1.37 | 825 | 850 |
| PASEF | 2 | 0.85 | 1.04 | 625 | 650 |
| PASEF | 2 | 0.64 | 0.85 | 425 | 450 |
| PASEF | 3 | 1.06 | 1.37 | 850 | 875 |
| PASEF | 3 | 0.87 | 1.06 | 650 | 675 |
| PASEF | 3 | 0.64 | 0.87 | 450 | 475 |
| PASEF | 4 | 1.09 | 1.37 | 875 | 900 |
| PASEF | 4 | 0.9 | 1.09 | 675 | 700 |
| PASEF | 4 | 0.64 | 0.9 | 475 | 500 |
| PASEF | 5 | 1.11 | 1.37 | 900 | 925 |
| PASEF | 5 | 0.92 | 1.11 | 700 | 725 |
| PASEF | 5 | 0.64 | 0.92 | 500 | 525 |
| PASEF | 6 | 1.13 | 1.37 | 925 | 950 |
| PASEF | 6 | 0.94 | 1.13 | 725 | 750 |
| PASEF | 6 | 0.64 | 0.94 | 525 | 550 |
| PASEF | 7 | 1.16 | 1.37 | 950 | 975 |
| PASEF | 7 | 0.97 | 1.16 | 750 | 775 |
| PASEF | 7 | 0.64 | 0.97 | 550 | 575 |
| PASEF | 8 | 1.18 | 1.37 | 975 | 1000 |
| PASEF | 8 | 0.99 | 1.18 | 775 | 800 |
| PASEF | 8 | 0.64 | 0.99 | 575 | 600 |

### Raw data analysis

Data were search against a human, reviewed, canonical FASTA downloaded from UniProt (20,376 entries, date: 19.04.2022). PROCAL only runs as well as the tissue data of the brain resourced were additionally searched against a PROCAL entry. DDA data were processed with MaxQuant (Version 1.6.17.0). Standard settings were used with 1% false discovery rate (FDR). Additionally, trypsin was set as protease allowing up to two missed cleavages, methionine oxidation and acetylation of N-termini were set as variable and carbamidomethylation of cysteine as fixed modification. Label-free quantification was activated with a minimal ratio count of 2. Match-between-runs was enabled for the tissue and body fluid data, but not the optimization and quality control data of the micro-flow timsTOF HT. The open search with FragPipe was performed using the “open” workflow (Version info: FragPipe version 19.1, MSFragger version 3.7, IonQuant version 1.8.10, Philosopher version 4.8.1). DIA data were analysed with Spectronaut17 by Biognosys AG (version: 17.1.221229.55965). Standard setting for library-based searches were used without cross run normalization and matching to FASTA was enabled. Custom-made libraries based on DDA runs acquired on the optimized micro-flow LC timsTOF HT setup were used (Single-shot plasma: 12,133 precursors of 556 protein groups; Single-shot CSF: 23,098 precursors of 2,227 protein groups; Deep fractionated HeLa: 299,276 precursors of 12,356 protein groups). Further data analysis was performed with Microsoft Excel, R and Python. Data visualization was performed with R, GraphPad 9.5.1 and BioRender (“Created with BioRender.com.”).

### Data processing

For each sample the medians were calculated from the subset of the proteins which were expressed in at least 70% of the samples. Afterwards, the intensities were corrected by multiplying to the correction factor which was obtained from dividing the average of medians by the median of that specific sample. To get an insight into the deferentially expressed proteins by pairwise comparisons across brain regions, the log transformed intensities values were subjected to the imputation of missing values using down shift normal distribution. (width = 0.3, downshift = 1.8). The resulting data for each protein were subjected to ANOVA followed by pairwise T-tests for multiple comparisons of independent groups (brain regions). The adjusted p-values were calculated by Benjamini/Hochberg method and the cut-off value of 0.05 was used to filter the proteins for each brain region. As reported in the supplementary table 1, a total number of 5750 proteins showed to be expressed significantly across different brain regions.

To establish reasonable criteria for calling fold changes significant, we considered a worst case scenario where all proteins have a CV of 40% (the highest variance in the brain region replicates analyzed in this study). Note that this is a very conservative assumption, as the vast majority of the proteins have much lower CVs. For this, we simulated the intensities for two brain regions, each with 3 biological replicates, from two normal distributions with a shared mean mu (randomly sampled from a normal distribution with mean: 1000.0, std: 200.0) and standard deviation mu*CV. This corresponds to the null hypothesis where there is no difference in means between the two brain regions. We repeated this for 10000 proteins, resulting in a distribution of fold changes calculated by dividing the means of the intensities in each of the two regions (Sup. Fig. 4.1D). This distribution of fold changes corresponds to an uncorrelated noncentral normal ratio (Hinkley et al., 1969), which has heavier tails than a log-normal distribution. This is a result of ratios from means drawn from opposite ends of the normal distribution. As expected, we observe that its analytical probability density function closely overlays with the simulated fold changes (Sup. Fig. 4.1E). We then applied numerical integration using the trapezoidal rule on the probability density distribution to obtain a cumulative density function (Sup. Fig. 4.1F). From this cumulative density function, we obtain that the two-tailed 95% confidence interval corresponds to a fold change of 2.02 (log2-fold-change=1.01). For the criterium of log2-fold-change>2, the corresponding two-tailed p-value is 1.6e-3.

In order to pinpoint proteins enriched in a brain region the classification scheme was adopted from (Sjöstedt *et al*., 2020). Class 1 encompasses proteins exclusively detected in one brain region within at least 2 biological replicates of that bran region and a maximum of one biological replicate in other brain regions. Class 2 includes proteins enriched in one brain region compared to all other brain regions in a pair-wise comparison by 4-fold (log2 difference of 2) considering the average of each brain region. Class 3 include group enriched proteins which reveal 4-fold enrichment in 2 to 7 brain regions compared the other brain regions. Class 4 encompasses brain region enhanced proteins which are 4-fold enriched compared to the average of the other brain regions.

To make the analysed data applicable to the front-end users, a shiny app was developed and dockerized in such a way that the user can explore the distribution of iBAQ or LFQ intensities of all identified proteins across different brain regions by searching for either gene name or UniProtID. In addition, an extra option was provided in the shiny app to make it possible for the selection and visualization of the annotated proteins according to their class categorizations or brain region enrichment.

To get an insight in to the similarity of the biological replicates obtained from different brain regions, the log transformed data for the most deferentially expressed proteins from the ANOVA results (as described earlier) were subjected to hierarchical cluster analysis using ward method and euclidean distance was selected as the metric before clustering. The resulting cluster-map is depicted in Supplementary Figure 4.2. Moreover, t-SNE plots were conducted to visualize proximity between the sample using the Rtsne package considering either all proteins in the region-resolved brain resource detected in a tissue sample or the average over the biological replicates of a brain region.

GO terms related to biological processes for human were retrieved using the R package GO.db and the pathways related to KEGG pathways were downloaded from the comparative toxicogenomics database (http://ctdbase.org/). In both cases the orthology mapping between ENTREZID and SYMBOL Ids was done using org.Hs.egGO. Afterwards, for each brain region, the proteins were primarily ranked according to their log2-fold change values and subsequently subjected to the fgsea package for fast gene set enrichment analysis. The pathways with positive normalized enrichment scores (NES) and p-adj < 0.05 were selected for further visualizations in Supplementary Figure 5.4. Functional annotation clustering with the DAVID tool was applied to compare proteins over- and underrepresented in the fresh tissue compared to formalin-fixed tissue (Huang da *et al*, 2009; Sherman *et al*, 2022).

## Data availability

Mass spectrometry raw data and search results performed with MaxQaunt or Spectronaut were deposited on Massive (Identifier: MSV000091835) (Choi *et al*, 2020). Moreover, a Shinyapp is available to the public (https://brain-region-atlas.proteomics.ls.tum.de/main_brainshinyapp/) summarizing the quantitative differences on iBAQ or LFQ level of the proteins within the brain resource enabling a direct comparison of a protein of interest between the 13 brain regions.

## Acknowledgments

This work was partially supported the Federal Ministry of Education and Research (Clinspect-M; FKZ161L0214A). We wish to thank the Bruker team involved in the project and all members of the Kuster group for their support. Study aim and brain region illustration was created with BioRender.com.

## Author contribution

JT, BK conceived the study. JT, EM, CK set up & optimized the micro-flow LC timsTOF system. JT, BK designed experiments. JT, SL performed experiments. JT, AS, MT performed bioinformatic data analysis. AS designed Shinyapp, JS, CD sampled tissue. JT, BK wrote the manuscript

## Conflict of interest

B.K. is a founder and shareholder of MSAID and OmicScouts. He has no operational role in either company. EM and CK are employees of Bruker Daltonics. The other authors declare no competing interests.

**Supplementary Figure 1.1**

A) iBAQ intensity distribution of proteins identified from a HeLa cell line digest in the presence or absence of 3% DMSO in LC solvents using a 30 min LC gradient and 2 µg injected digest (N=3).

B) Distribution of ion mobility elution times [ms] of peptides detected in A).

C) Number of identified peptides measured using a 30 min LC gradient as function of the amount of HeLa peptides injected using different electrospray emitters of 50, 80 or 100 µm inner diameter (ID) (N=3).

D) Distribution of the chromatographic peak width distribution (full width, FW) of peptides in panel C).

E) Bar graph showing the number of unique peptides and protein groups identified by a 30 min HeLa run (2 µg) comparing different MS2 scheduling thresholds (N=3) E)

F) and MS2 scheduling speeds F). Time spend per precursor for MS2 scanning including quadrupole switching time was reduced from standard (Std) of 4.4 ms to fast settings of 3.2 ms.

G) Number of scheduled precursors for fragmentation per 100 ms ramp using the fast or standard (Std) method during a 30 min HeLa gradient run (2 µg).

H) Bar graph showing the number of protein groups and unique peptides identified within a 30 min HeLa run (2 µg) as a function of different collision energy settings between 45 to 27 and 61 to 29 (N=3).

I) Table summarizing the data shown in panel H).

**Supplementary Figure 1.2**

A) Number of identified peptides (upper panels) and proteins (lower panels) by micro-flow LC-MS/MS (30 min LC gradient) using a timsTOF Pro2 (red) or timsTOF HT (blue) mass spectrometer. HeLa digest dilutions were analysed in triplicates using between 0-10 µg (in ascending order) (N=3).

B) Absolute (upper panels) and relative (bottom panels) number of identified fully tryptic or semi-tryptic peptides as a function of the amount of HeLa digest injected and analysed by micro-flow LC-MS/MS (30 min LC gradient) using a timsTOF Pro2 (red) or timsTOF HT (blue) mass spectrometer.

**Supplementary Figure 2.1**

A) Number of identified peptides (top and middle panels) analysed by data dependent acquisition (DDA, light green) or data independent acquisition (DIA, dark green) analysed by different LC gradients and as a function of the amount of HeLa digest injected (N=3). The lower panels show the number of identified proteins from the panels above in direct comparison.

B) Cumulative density plot of the number of proteins as a function of quantitative precision (coefficient of variation (CV)) using protein LFQ intensities on protein level (2 µg HeLa peptides, 30 min LC gradients, N=5).

C) Same data as in panel A) but showing the number of protein identifications as a function of the amount of HeLa digest injected and analysed by different LC gradients in DDA and DIA mode.

**Supplementary Figure 2.2**

A) Left panel: Number of identified peptides from undepleted human plasma digests analysed by data dependent acquisition (DDA) and different LC gradients and as a function of the amount of sample injected. Right panel: Number of identified peptides from undepleted human plasma digest (5 human individuals) injecting 5 µg digest and analysed using a 30 min LC gradient and by DDA and DIA.

B) Same as panel A) but for human CSF.

C) Dose–response binding curves of targets of the HDAC inhibitor TSA obtained by competition binding assays to HDAC beads and analysed by micro-flow timsTOF LC-MS/MS (light colors) or nano-flow Orbitrap MS/MS (dark colors). The latter data was reproduced from (Lechner *et al*., 2022).

D) Correlation analysis of -log 10 EC 50 values of dose-response curves shown in panel C). R, Pearson correlation coefficient.

**Supplementary Figure 3**

A) Correlation analysis of the mean retention time of PROCAL peptides analysed alone (N=10) or spiked into tissue samples (N=1968, 30 min gradient). Peptides detected in at least 75% of the spike-in cases were considered.

B) Summary of comparing protein identifications from fresh frozen to formalin-fixed regarding intensity and overlap indicating differences between quantiles. Quantile 1 represents the 25% most abundant proteins.

C) Density scatter plot showing the log10 iBAQ intensity of proteins in fresh sample vs. the ratio of fresh and fixed cortical samples. Dotted lines mark the 10% of proteins with the strongest deviation between fresh and FFPE samples.

D) Results of functional annotation clustering using DAVID of the top and bottom 10% of proteins overrepresented/ underrepresented in fresh samples. The circle area is proportional to the functional annotation score.

**Supplementary Figure 4.1**

A) Overlap of protein identifications between any two or more brain regions (proteins were required to have been detected in at least two biological replicates).

B) Same as panel A) excluding proteins shared between all brain regions.

C) Bar graphs indicating the absolute and relative number of proteins in different enrichment classes for each brain region.

D) Simulated data of the fold-change between the mean intensities of the biological replicates of proteins in two brain regions with the same mean and CV= 40% (the highest variance in the brain region replicates analyzed in this study) N=3. For a given CV value, the fold change distribution is independent of the intensity values. The red dashed lines represent the 95% confidence interval as determined by the distribution in panel E, whereas the green dashed lines represent the cutoff of log2 fold-change of 2. See methods for details.

E) Histogram of the fold changes from panel D, overlaid with the probability density function of the corresponding uncorrelated noncentral normal ratio distribution for CV= 40%. The analytical distribution closely overlays the histogram.

F) Cumulative density function corresponding to the probability density function of panel E. The 95% confidence interval corresponds to a fold change of 2.02 (log2 fold-change= 1.01), whereas the log2 fold-change= 2 corresponds to a p-value of 1.6e-3.

**Supplementary Figure 4.2**

Scatter plot indicating the correlation of the proteomic brain region data acquired in the scope of this project compared to RNA data reported human protein atlas (HPA) (Sjöstedt *et al*., 2020). The five regions included in both omic studies are shown.

**Supplementary Figure 4.3**

A) tSNE plot illustrating proximity of proteomes between brain regions (biological replicates were averaged and are shown as single dots per brain region). Basal ganglia (NAC, CN, PUT) and midbrain (NR, SN) samples are highlighted with a dotted grey line.

B) Comparison of protein and mRNA levels of example proteins in five brain regions. The mRNA data was taken from the human protein atlas (HPA) (Sjöstedt *et al*., 2020). Each panel highlights examples for each brain regions in which the proteins shown are statistically significantly higher expressed than in other brain regions.

C) Examples for protein expression differences across the13 brain regions.

D) Swarm plot showing Class2 proteins sorted by LFQ intensity.

**Supplementary Figure 4.4**

Hierarchical clustering and heat map visualization of all Anova-positive proteins indicating significantly different abundance levels of a proteins between brain regions.

**Supplementary Figure 5.1**

A) Protein abundance distribution of all proteins in the brain proteome atlas (blue) and the region-enriched proteins (yellow).

B) Protein abundance ratios of all astrocyte-, microglia- oligodendrocyte- and neuron-enhanced proteins (according to HPA) in a specific brain region plotted over the average of all other brain regions (dotted line).

C) Protein abundance profiles of three neuropeptides that exhibit enrichment in a specific brain region.

**Supplementary Figures 5.2 + 5.3**

Mean (N=3) log2 iBAQ intensities of the top 20 most abundant proteins in a particular brain region plotted over the difference to the mean of all other brain regions.

**Supplementary Figure 5.4**

KEGG and GO-enrichment (biological processes) of the proteins included in the region resolved resource. Proteins are ranked according to their fold change difference to the average of all brain regions. The size of the dots indicates the number of enriched proteins (higher than 4-fold) linked to the enrichment term and the colour indicates the activation score.

